# CryoVirusDB: A Labeled Cryo-EM Image Dataset for AI-Driven Virus Particle Picking

**DOI:** 10.1101/2023.12.25.573312

**Authors:** Rajan Gyawali, Ashwin Dhakal, Liguo Wang, Jianlin Cheng

## Abstract

With the advancements in instrumentation, image processing algorithms, and computational capabilities, single-particle electron cryo-microscopy (cryo-EM) has achieved nearly atomic resolution in determining the 3D structures of viruses. The virus structures play a crucial role in studying their biological function and advancing the development of antiviral vaccines and treatments. Despite the effectiveness of artificial intelligence (AI) in general image processing, its development for identifying and extracting virus particles from cryo-EM micrographs (images) has been hindered by the lack of manually labelled high-quality datasets. To fill the gap, we introduce CryoVirusDB, a labeled dataset containing the coordinates of expert-picked virus particles in cryo-EM micrographs. CryoVirusDB comprises 9,941 micrographs of 9 different viruses along with the coordinates of 339,398 labeled virus particles. It can be used to train and test AI and machine learning (e.g., deep learning) methods to accurately identify virus particles in cryo-EM micrographs for building atomic 3D structural models for viruses.

## Background & Summary

Cryo-electron microscopy (cryo-EM) is a method of capturing 2D images of biological molecules and assemblies at extremely low (cryogenic) temperatures. Advancements in both instrumentation and computational methodologies have established cryo-EM as an essential tool for interrogating the structures and dynamics of biological macromolecular complexes including large virus particles [1]. Single particle experiment and data analysis in cryo-EM involves flash-freezing biological specimens, collecting micrographs of individual particles in an electron microscope (**Figure 1A**), followed by picking and extracting particle images (**Figure 1B**), applying image processing for correction and alignment (**Figure 1C**), and performing three-dimensional (3D) reconstruction of macromolecular complexes (**Figure 1D**) [1], [2].

**Figure 1:**
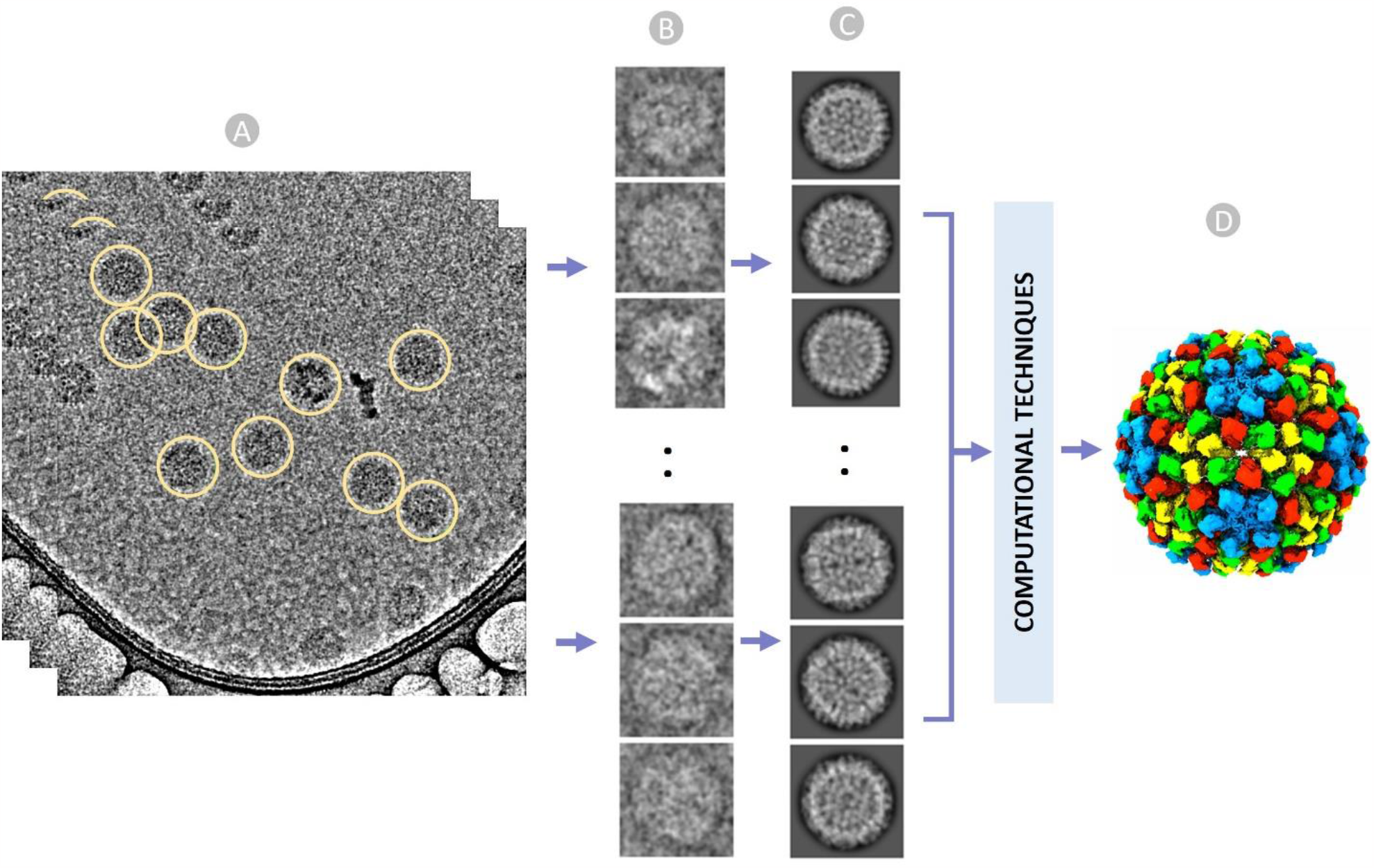
An overview of cryo-EM single particle analysis from particle selection to 3D reconstruction of virus. (A) Stack of ideal micrographs where the true virus particles are picked (encircled yellow), (B) Extracted virus particles from micrographs with fixed box size. (C) Multiple 2D classes to facilitate stack cleaning and the removal of false particles. (D) Reconstructed 3D structure of the virus from 2D images using a series of computational techniques.

In the realm of virology, cryo-EM has been instrumental in studying and determining the 3D structures and morphology of various viruses such as Polio, Ebola, HIV, and Corona viruses [3] [4]. Particularly during the COVID-19 pandemic, cryo-EM played a pivotal role in understanding the intricate structure of the SARS-CoV-2 spike protein [5] [6]. This knowledge has facilitated the development of highly effective vaccines. For instance, scientists have been able to design immunogens that mimic the spike protein’s shape, eliciting targeted immune responses [7]–[9]. Moreover, cryo-EM has revolutionized epitope mapping, enabling the identification of specific binding sites [10] and facilitating the exploration of antibody mutations for the rapid discovery and development of precise vaccines and antiviral treatments [11] [12] [13].

To achieve high-resolution 3D reconstructions of virus structures, the initial step of accurately recognizing and extracting virus particles from 2D image projections (micrographs) is crucial. Currently, three virus particle picking approaches are employed: manual virus particle picking, template-based picking, and AI-based picking. Manual picking is laborious and time-consuming, requiring specialized expertise for precise identification, which cannot be used by regular users. Challenges in the manual picking arise from low single-to-noise ratios, low particle contrast, and the unpredictability of individual particle appearances due to orientation variations. Template-based virus particle picking requires experts to pick some initial particles as templates for software tools to search for more particles, which suffers from the presence of ice contamination, radiation damaged particles, carbon areas, and overlapping aggregated particles in micrographs. AI-based particle picking [14] [15] [16] has the best potential to automate the process and overcome the problems of the manual picking and template-based matching, but the development of sophisticated AI-based virus particle picking methods is largely hindered by the lack of high-quality labelled training and test data of virus particles.

To harness the power of cutting-edge AI technologies in automatic virus particle recognition and picking, we created a comprehensive and expert-labelled dataset – CroVirusDB -[17] in this work. This open-access dataset aims to expedite the development of automated virus particle picking workflows, and ultimately advance the research of viruses and the design of therapeutic interventions. CryoVirusDB includes 9,941 micrographs of 9 distinct viruses and the coordinates of 339,398 virus particles picked in them. The statistics of CroVirusDB is reported in **Table 1 Table 2.**

**Table 1:** The statistics of micrographs and particles of 9 viruses in CryoVirusDB.

| SN | EMPAIR ID | Virus Type | Number of Micrographs | Micrograph size | Particle Diameter (px) | Number of True Virus Particles |
| --- | --- | --- | --- | --- | --- | --- |
| 1 | 10192 [18] | Feline calicivirus | 1000 | (4096, 4096) | 470 | 9660 |
| 2 | 11060 [19] | Nudaurelia capensis omega virus | 1276 | (4096, 4096) | 516 | 11916 |
| 3 | 10203 [20] | Macrobrachium rosenbergii nodavirus | 1000 | (3838, 3710) | 377 | 16601 |
| 4 | 10033 [21] | Human parechovirus 3 | 1000 | (4096, 4096) | 350 | 55732 |
| 5 | 10652 [22] | Coxsackievirus | 1127 | (3838, 3710) | 374 | 11144 |
| 6 | 10341 [23] | Bovine enterovirus | 1274 | (4096, 4096) | 376 | 22694 |
| 7 | 10193 [18] | Feline calicivirus | 1000 | (4096, 4096) | 516 | 96126 |
| 8 | 10205 [24] | Cowpea mosaic virus | 1000 | (4096, 4096) | 310 | 81037 |
| 9 | 10555 [25] | Nudaurelia capensis omega virus | 1264 | (4096, 4096) | 564 | 34488 |
|  |  | **Total** | **9941** |  |  | **339,398** |

**Table 2:** Metrics of EM data acquisition and grid preparation utilized in importing micrographs for virus particle picking.

| SN | EMPAIR ID | Micrograph Format | Pixel Spacing (Å) | Accl Voltage (kV) | Spherical Aberration (mm) | Electron Dose (e/A<sup>2</sup>) | Defocus range (µm) | Microscope | Detector |
| --- | --- | --- | --- | --- | --- | --- | --- | --- | --- |
| 1 | 10192 | mrc | 1.065 | 300 | 2.7 | 63 | NA | FEI TITAN KRIOS | FEI FALCON III (4k x 4k) |
| 2 | 11060 | mrc | 1.065 | 300 | 2.7 | 46 | 0.70 µm - 2.2 µm | FEI TITAN KRIOS | FEI FALCON III (4k x 4k) |
| 3 | 10203 | mrc | 1.06 | 300 | 2.7 | 36 | 1.0 µm - 2.5 µm | FEI TITAN KRIOS | GATAN K2 SUMMIT (4k x 4k) |
| 4 | 10033 | mrc | 1.14 | 300 | 2.7 | 36 | 0.42 µm - 2.34 µm | FEI TITAN KRIOS | FEI FALCON II (4k x 4k) |
| 5 | 10652 | mrc | 1.06 | 300 | 2.7 | 40 | 0.6 µm - 3.0 µm | TFS TALOS F200C | FEI FALCON III (4k x 4k) |
| 6 | 10341 | mrc | 1.065 | 300 | 2.7 | 49.5 | 0.75 µm - 3.5 µm | FEI TITAN KRIOS | FEI FALCON III (4k x 4k) |
| 7 | 10193 | mrc | 1.065 | 300 | 2.7 | 63 | NA | FEI TITAN KRIOS | FEI FALCON III (4k x 4k) |
| 8 | 10205 | mrc | 1.065 | 300 | 2.7 | 67.5 | NA | FEI TITAN KRIOS | GATAN K2 SUMMIT (4k x 4k) |
| 9 | 10555 | mrc | 1.0651 | 300 | 2.7 | 72 | 0.70 µm - 2.7 µm | FEI TITAN KRIOS | FEI FALCON III (4k x 4k) |

## Methods

### 1. Raw Data Acquisition and Preprocessing

The metadata and cryo-EM virus micrographs from the EMPIAR web portal [26] were fetched using Python API and FTP scripts. The comprehensive metadata encompasses the EMPIAR ID for each cryo-EM dataset of a virus along with the corresponding identifiers such as Electron Microscopy Data Bank (EMDB) ID and Protein Data Bank (PDB) ID. Additionally, the dataset size, resolution, total number of micrographs, image specifications (size and type), pixel spacing, micrograph file extension, gain/motion correction file extension (if any), FTP and Globus paths for micrograph/gain files, and relevant publication information are meticulously recorded.

To ensure dataset diversity, we selected 9 representative EMPIAR virus datasets that encompassed a broad range of particle sizes, shapes, density distributions, noise levels, and variations in ice thickness and carbon areas to create CryoVirusDB. The datasets include viruses from different categories, such as Omage virus, Cowpea Mosaic virus, Feline calicivirus, and Human parechovirus, providing a comprehensive representation of the virus space.

For each EMPIAR virus dataset, we imported its raw micrographs. A meticulous analysis of the EM data acquisition descriptions and grid preparation details for each dataset was undertaken to gather essential information such as raw pixel size (Å), acceleration voltage (kV), spherical aberration (mm), and total exposure dose (e/Å 2) associated with the micrographs in the respective dataset as shown in **Table 2**.

### 2. Motion Correction and Patch based CTF Estimation of Micrographs

In our study, we used motion corrected micrographs as the starting point in CryoSPARC [27] for patch-based Contrast Transfer Function (CTF) estimation. Since CTF functions can vary substantially among micrographs and cannot be precisely predefined, accurately identifying CTF parameters for each micrograph is crucial. This precision is necessary for proper corrections and achieving high-resolution 3D reconstructions. Two stages (estimating the CTF and correcting it) are applied to the CTF analysis.

We employ the patch-based Contrast Transfer Function (CTF) to generate output micrographs containing information about their average defocus and the defocus landscape. Upon particle extraction, this information is automatically utilized to allocate a local defocus value to each particle based on its position in the landscape. The one-dimensional search across defocus values for a micrograph is shown in **Figure 2A**.

**Figure 2:**
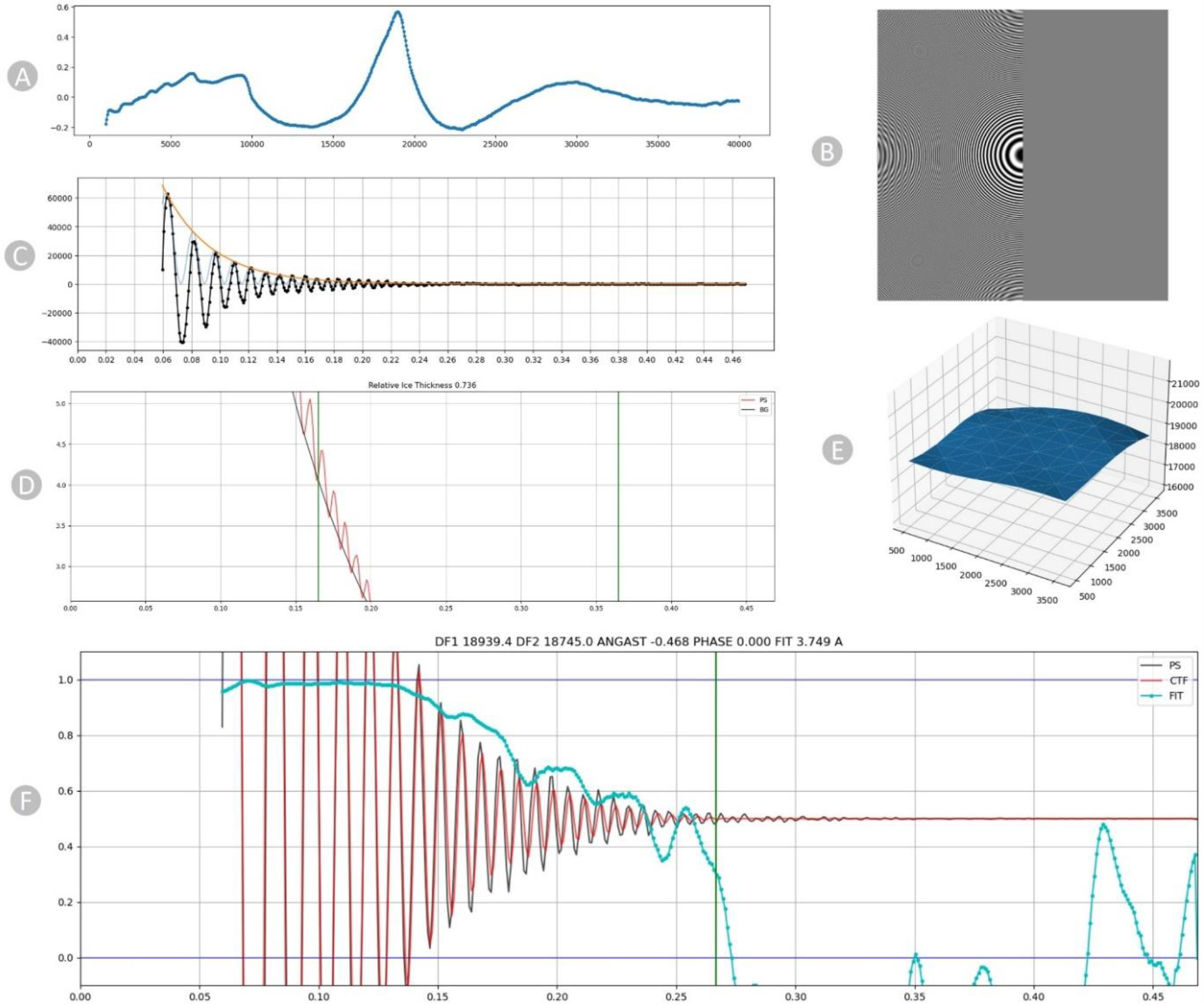
Diagnostic plots of CTF for EMPAIR 11060. (A) 1D search over varying defocus values. (B) Thon rings visible in the Fourier transform. (C) Fitted envelope function diagnostic plot. (D) Power Spectrum showing the relative ice thickness. (E) 2D Patch result. (F) CTF fit plot. The frequency, measured in inverse angstroms (Å^-1^), is represented on the X-axis, while the correlation metric between the power spectrum (PS) and CTF value is shown on the Y-axis. The black line corresponds to the observed experimental power spectrum, the red line represents the calculated CTF, and the cyan line indicates the cross-correlation (fit).

The distinctive characteristic of the Contrast Transfer Function (CTF) is its oscillating pattern, easily observable as Thon rings in the power spectra of images (**Figure 2B**). Thon rings exhibit more frequent oscillations with larger defocus values and fewer oscillations with smaller defocus values. This connection between defocus and Thon rings forms the foundation for both manual and automated methods of fitting the CTF.

The plot in **Figure 2C** serves primarily as a verification for the successful execution of background subtraction and envelope function fitting. The X-axis represents frequency in inverse angstroms. The radially averaged power spectrum is depicted in black, where high values correspond to the bright portions of the Thon rings and low values to the dark regions. The orange curve represents the envelope function, aiming to model the expected falloff of Thon rings up to the Nyquist resolution [28], accounting for aberrations. Lastly, the fitted Contrast Transfer Function (CTF), scaled by the envelope function, is presented in blue. This oscillating plot is crucial for confirming the proper execution of background subtraction and envelope fitting procedures. In the plot in **Figure 2D**, we assess the background strength (depicted by the black line) within the area where thicker ice leads to an augmented background referred to as relative ice thickness.

The 3D surface plot (**Figure 2E**) shows the local defocus estimated throughout the micrograph. The surface plot in blue illustrates the defocus that has been fitted for each position along the micrograph. The x- and y-coordinates align with the micrograph’s coordinates, while the z-coordinate represents the defocus values. The X, Y, and Z axes are all expressed in Angstrom units. The Contrast Transfer Function fit plot, illustrated in **Figure 2F**, depicts the alignment between the simulated and observed Thon rings in the micrograph, accounting for variations in defocus and astigmatism. The cyan curve indicates the cross-correlation fit level. The CTF fit resolution (3.749 angstroms) is the resolution at which this value drops below a threshold. The vertical green line in the plot signifies the frequency at which the fit deviates from cross-correlation threshold of 0.3, indicating a successful fit.

### 3. Manual Particle Picking and 2D Class Formation

Following the CTF estimation, we manually identified and selected true virus particles interactively from aligned and motion-corrected micrographs with the aim of generating some particle templates. We specify the particle diameter based on the virus particles’ size and shape. Picking particles directly from raw noisy micrographs is challenging (**Figure 3A)**. So we adjusted the ‘Contrast Intensity Override’ using low pass filter while inspecting micrographs to achieve the most distinct view for particle selection (**Figure 3B)**. Additionally, we employed a visual guide to encircle virus particles (**Figure 3C)**, ensuring that the chosen particles are well-centered for improved results in subsequent 2D alignment steps.

**Figure 3:**
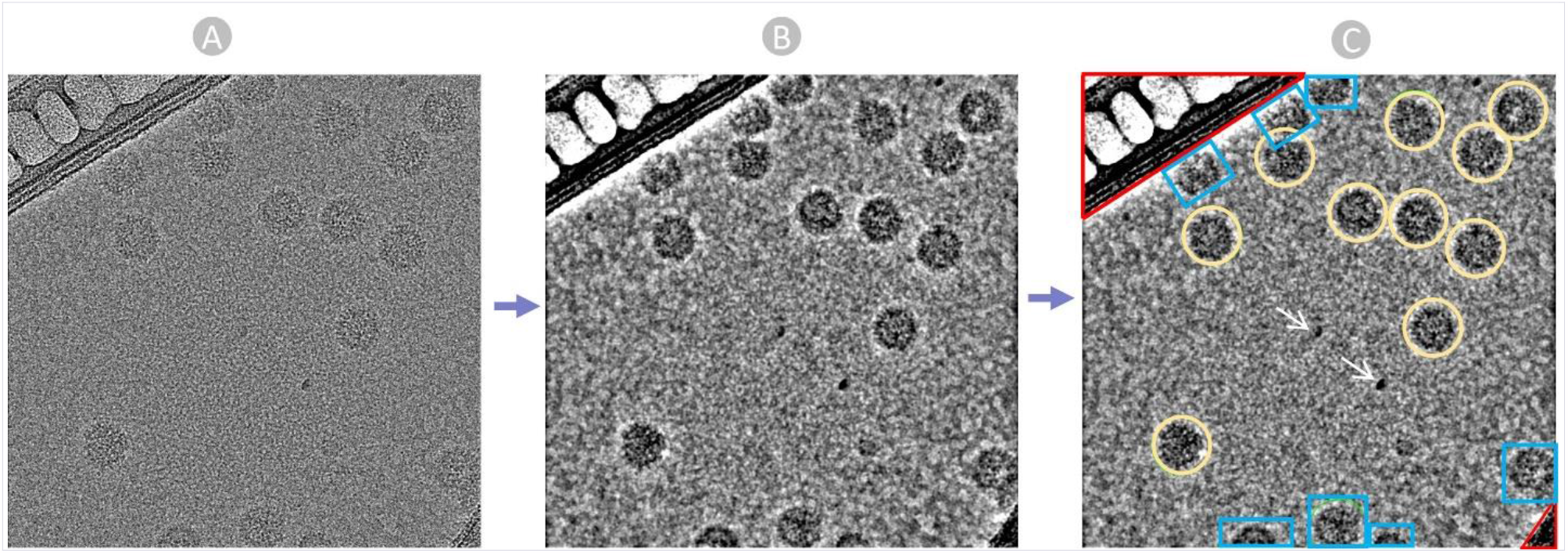
The manual Picking Process. (A) Raw micrograph obtained from EMPIAR. (B) Preprocessed Micrograph with Low Pass filter: 28 A to ease particle recognition and picking. (C) Manually picked true virus particle encircled in yellow, carbon region colored in Red, ice patches and artifacts pointed by white arrows, and cut particles in edges colored in blue.

Manually selecting particles from raw micrographs with smaller defocus values proves to be quite challenging. To create a comprehensive set of ground-truth templates covering a broad range of defocus values, we manually picked particles from numerous micrographs exhibiting diverse defocus and CTF fit values. Given the time-intensive nature of manual picking, we chose a small subset of micrographs (around 15% of the micrographs) specifically for generating templates. The detailed information about the manually picked particles and the micrographs considered for the manual picking can be found in **Supplementary Table S1**.

After manually picking the virus particles, the coordinates of particle centers and the designated box size are used to extract particles from the original micrographs. During this process, the box size is defined to provide ample padding, usually ranging from 25% to 50% extra space around the particles. The manually selected particles undergo a 2D classification step, where we categorized and chose the most favorable classes. This classification step organized particles into distinct 2D classes, streamlining the cleaning of the particle stack and removal of undesirable particles. Finally, we assessed the quality of the particles and eliminated classes containing unwanted particles. The remaining particle classes are used by the template-based picking for the identification of high-quality particle classes.

### 4. Template-based Picking

After exporting the optimal particle classes, we employed templates created in the ‘2D Class Formation’ step in CryoSPARC. We followed an iterative approach, wherein the output from ‘template-based picking and inspection’ is once again utilized in the ‘2D Class Formation’ step to select only the high quality 2D particles discarding the false positives. This cycle was repeated until we obtained high-resolution particles that encompass all possible viewing directions of the virus particle.

Using CryoSPARC’s Template Picker job, we employed the high-resolution templates to precisely pick virus particles that align with the geometry of the target structure. We set specific constraints, such as the Particle diameter in angstrom and a minimum distance between the particles for generating templates based on the SK97 sampling algorithm [29].

### 5. Manual Particle Inspection and Extraction

The acquired particles above underwent the manual inspection, in which we scrutinized and refined the picked particles using different thresholds. We fine-tuned parameters such as the lowpass filter, normalized cross-correlation (NCC), and power threshold (**Figure 4A**) to eliminate false positives. The 2D colored histogram plots were employed to carefully analyze the median pick scores of micrographs against defocus, aiding in the extraction of coordinates for high-quality virus particles as depicted in **Figure 4B**.

**Figure 4:**
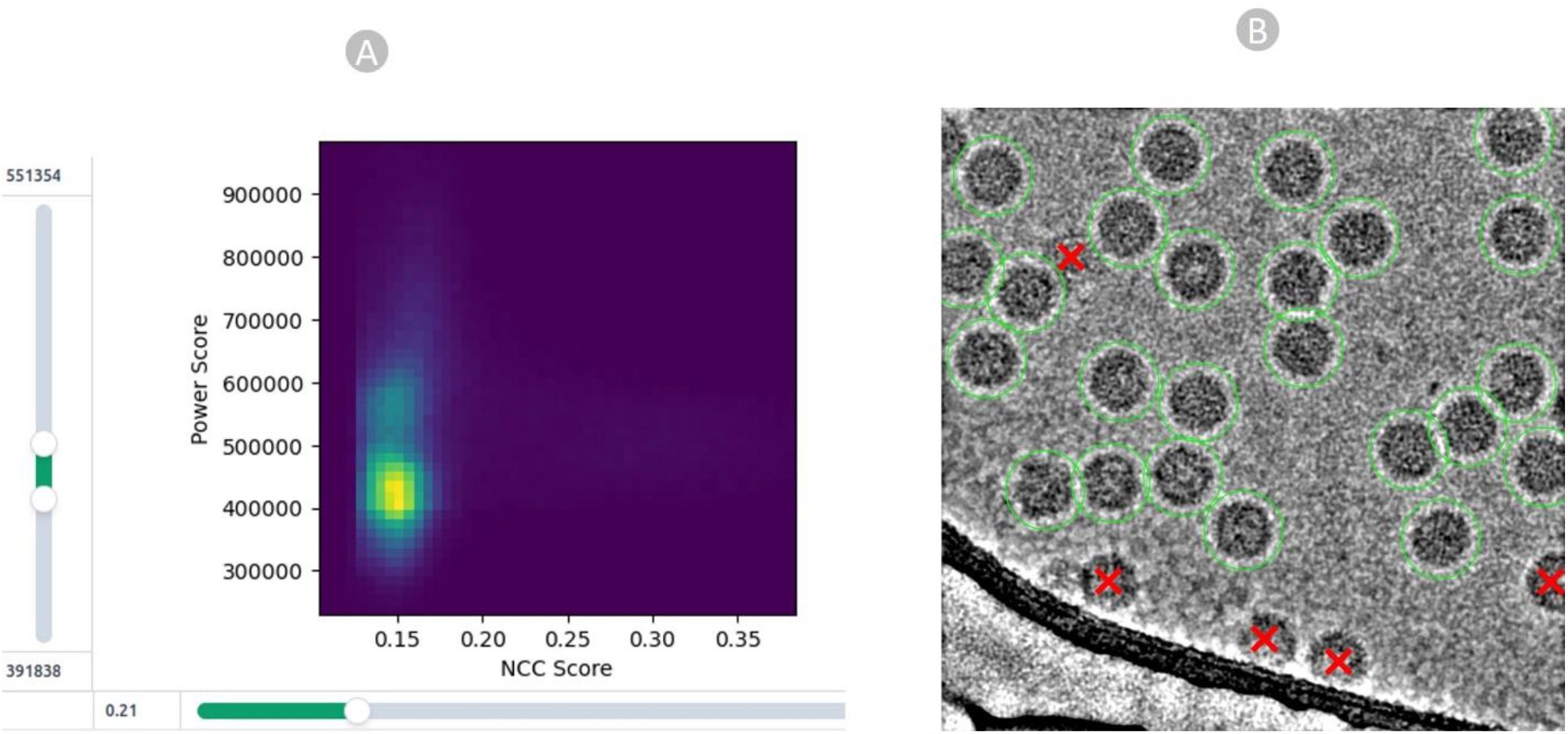
Particle Quality inspection. (A) particle filtration achieved by manipulating the NCC score on the X-axis and local power score on the Y-axis for EMPIAR 11060. (B) high quality true virus particles (depicted by green circles) chosen through the template-based picking process and eliminated radiation-damaged, cut, and false positive particles, represented by red crossed markers.

Ultimately, we applied a 2D Classification step to perform a final examination of the selected particles. The Select 2D job categorized particles into various 2D classes (usually 50 in our case), aiding in stack cleaning and the elimination of undesired particles. This process is valuable not only for assessing particle quality before entering the 3D reconstruction phase but also for qualitatively exploring the distribution of views within the dataset. Following 2D Classification, certain classes are identified as “junk” classes, representing non-particle images, ice crystals, or instances of two particles being conjoined. Consequently, we filtered out the particles associated with these “junk” classes from the picked particles. More information about the overall intermediate metadata and the final set of true virus particles can be found in **Supplementary Table S1**.

These final true particles are exported in the form of particle stacks, star files and csv files, which include a lot of information about the particles in micrographs like: X-coordinate, Y-coordinate, Angle-Psi, Origin X (Ang), Origin Y (Ang), Defocus U, Defocus V, Defocus Angle, Phase Shift, CTF B Factor, Optics Group, and Class Number.

## Data Records

CryoVirusDB includes 9 virus subsets (each including approximately 1200 cryo-EM micrographs) along with the labelled coordinates of the virus particles in the micrographs. The total size of the CryoVirusDB database is 634 GB. The organizational structure of the directories of CryoVirusDB is depicted in **Figure 5**.

**Figure 5:**
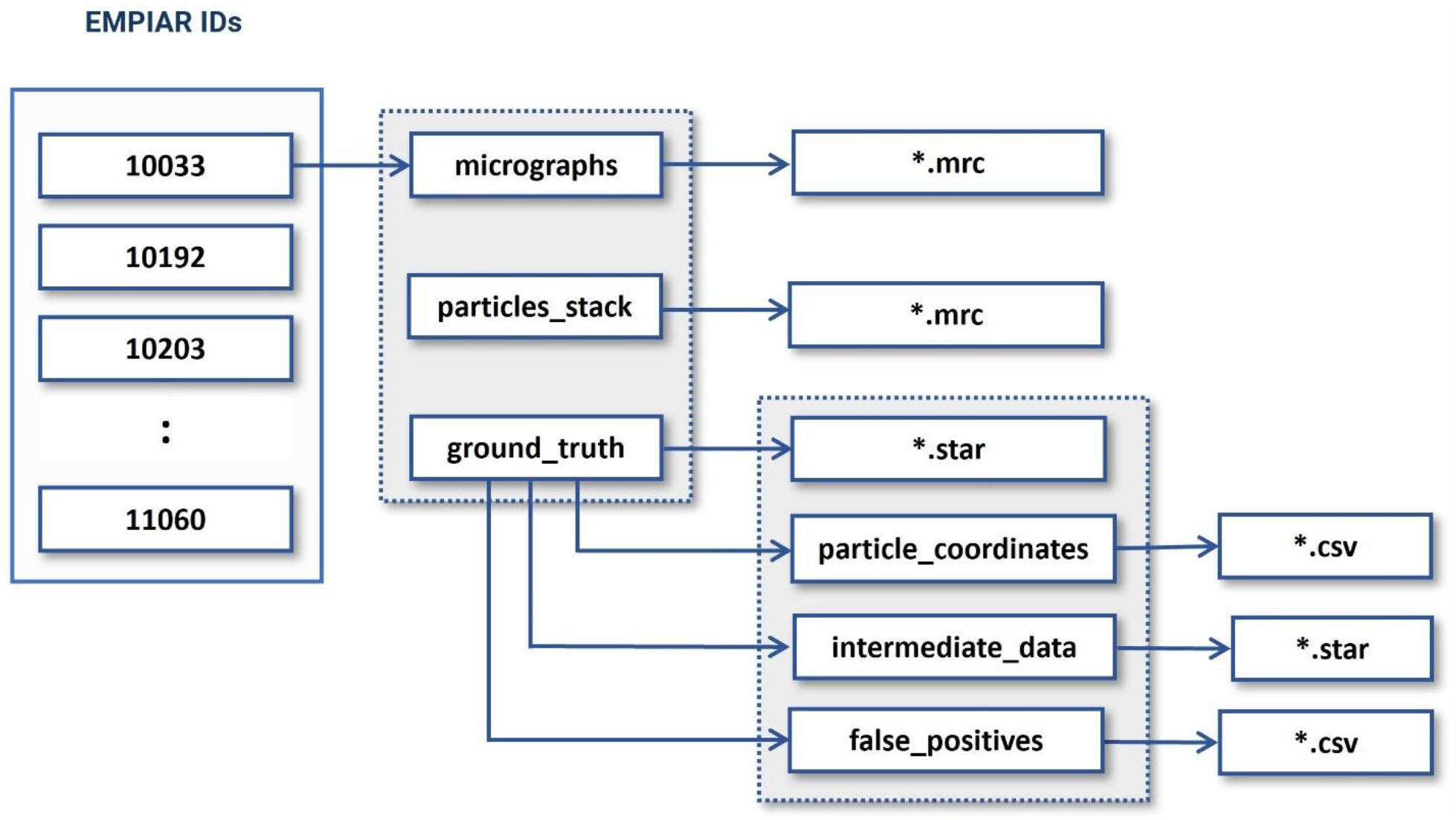
The directory structure of CryoVirusDB. The numbers in the blocks on the left side are the respective EMPIAR IDs.

### 1. Motion Corrected Micrographs

These are the two-dimensional images captured by the microscope during the imaging process. All the micrographs in CryoVirusDB are stored in .*mrc* image format. Each sub-dataset (named with EMPIAR ID) in CryoVirusDB contains around 1200 micrographs.

### 2. Virus Particle Stack

The particle stack consists of .*mrc* files, each named after the corresponding micrograph’s filename, containing ground truth virus particles. These files form a three-dimensional grid of voxels, where each voxel value corresponds to electron density, essentially forming a stack of 2D images. To view and inspect the particle stacks, one can use EMAN2 [30] or UCSF Chimera [31] / ChimeraX [32] .

### 3. Ground Truth Labels (Coordinates)

The ground truth directory includes particle coordinates (in .csv format), false positives (in .csv format), intermediate data (in .star file), and the collective star file of all ground truth particles. The false positives contain viruses like particles that are actually ice contaminations, aggregates, radiation damaged particles, and false particles over carbon regions.

## Technical Validation

### 1. 2D Particle Class Validation

We compared our picked virus particles with a popular AI-based particle picking method, Topaz [33], considering factors such as the total number of classes, number of picked particles, 2D resolution, and visual orientation. Our manually picked particles have a better 2D class resolution than Topaz. It’s noteworthy that a higher particle count alone does not ensure higher resolution. Instead, selecting a substantial number of high-quality particles across a broad angular distribution is crucial for achieving both high 2D and 3D resolution. The 2D class comparison for two databases: EMPIAR 10205 and EMPIAR 10193 (each containing 1000 micrographs) are shown in **Table 3** and **Figure 6**. In both cases, Topaz picked many more particles than CryoVirusDB but it had a worse 2D class resolution.

**Table 3:** 2D classification result comparison for EMPIAR 10205 and EMPIAR 10193.

| EMPIAR 10205 |  |  |
| --- | --- | --- |
|  | 2D Particle Class Statistics<br>(Topaz) | 2D Particle Class Statistics<br>(CryoVirusDB) |
| Number of Picked Particles | 155,953 | 81,037 |
| Weighted Average Resolution of 2D classes<br>(N=50) | 9.41 Å | 6.59 Å |
| Weighted Average Resolution of 2D classes<br>(N=10) | 13.42 Å | 10.96 Å |

| EMPAIR 10193 |  |  |
| --- | --- | --- |
|  | 2D Particle Class Statistics<br>(Topaz) | 2D Particle Class Statistics<br>(CryoVirusDB) |
| Number of Picked Particles | 239,852 | 96,126 |
| Weighted Average Resolution of 2D classes<br>(N=50) | 18.52 Å | 15.02 Å |
| Weighted Average Resolution of 2D classes<br>(N=10) | 23.68 Å | 21.72 Å |

**Figure 6:**
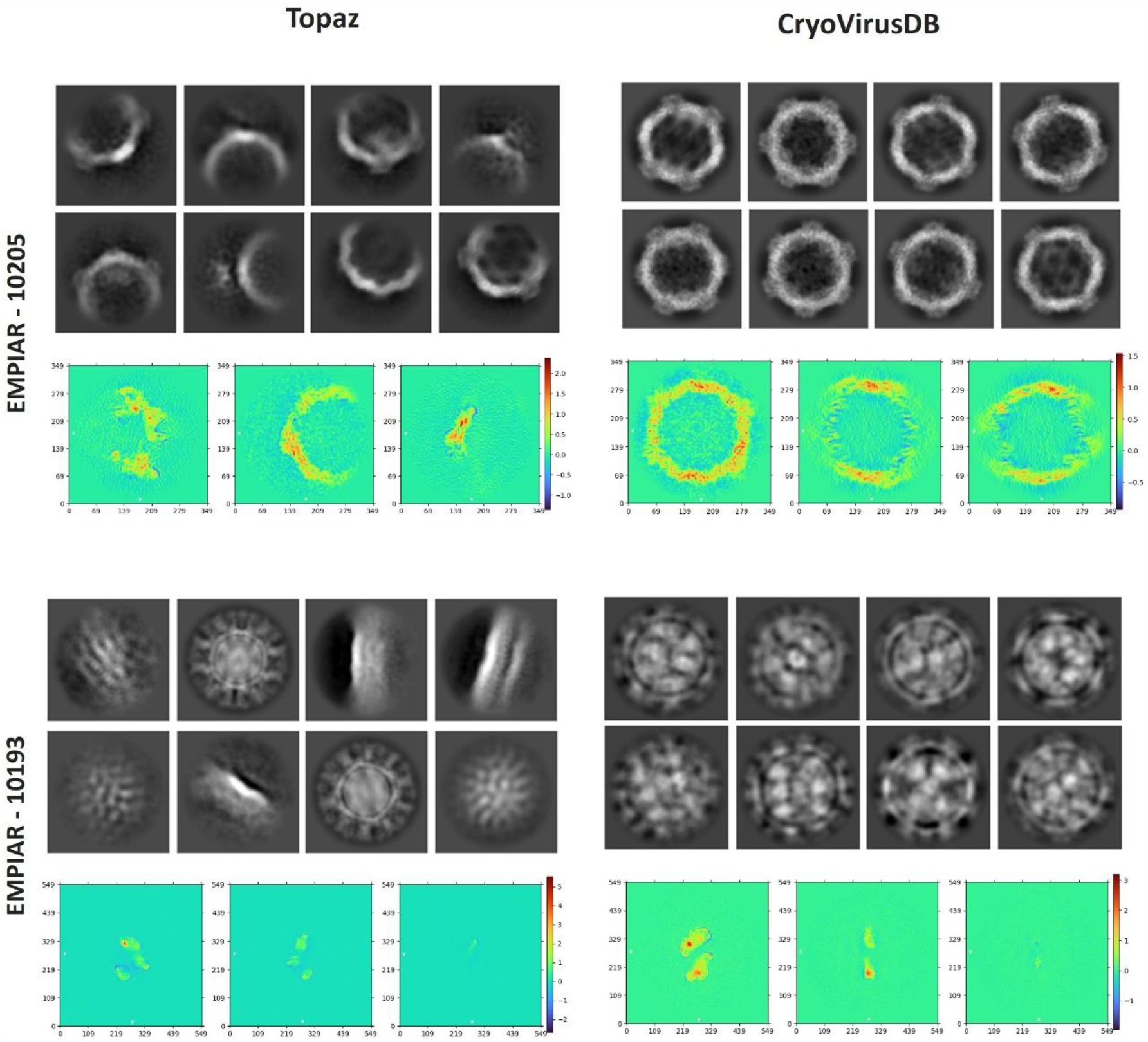
The comparison of 2D particle classification and density projections from the intermediate output of the ab initio reconstruction phase for EMPIAR 10205 and EMPIAR 10193. For each EMPIAR ID, the 2D classes are visualized at the top and the density projects are visualized at the bottom. The color scheme in the heatmap corresponds to the scalar density values at each voxel.

We also assessed the density projections derived from the intermediate output during the ab initio reconstruction phase, as depicted at the bottom of each block in **Figure 6**. The plot illustrates the integrated density values along the perpendicular direction to that plane. The heatmap’s color scheme represents scalar density values at each voxel, with the intensity of color indicating the magnitude of density. This indicates the high quality of the virus particles in CryoVirusDB.

### 2. 3D Density Map Validation

We reconstructed 3D density maps from the particles in CryoVirusDB and from those picked by Topaz for two datasets: EMPIAR 10205 and EMPIAR 10193, each comprising 1000 micrographs. The ab-initio density map reconstruction and homogenous refinement were carried out in CryoSPARC using the generated star files that included the selected particles. To ensure an unbiased evaluation, we repeated the ab-initio 3D reconstruction experiment with three distinct random seeds for each method.

**Figure 7** presents a comparison of the resolution and distribution direction of the reconstructed 3D density maps. The Fourier Shell Correlation (FSC) plots include a ‘loose mask’ curve that utilizes an automatically generated mask with a 15 Å falloff, and a ‘tight mask’ curve that employs an auto-generated mask with a falloff of 6 Å for all FSC plots.

**Figure 7:**
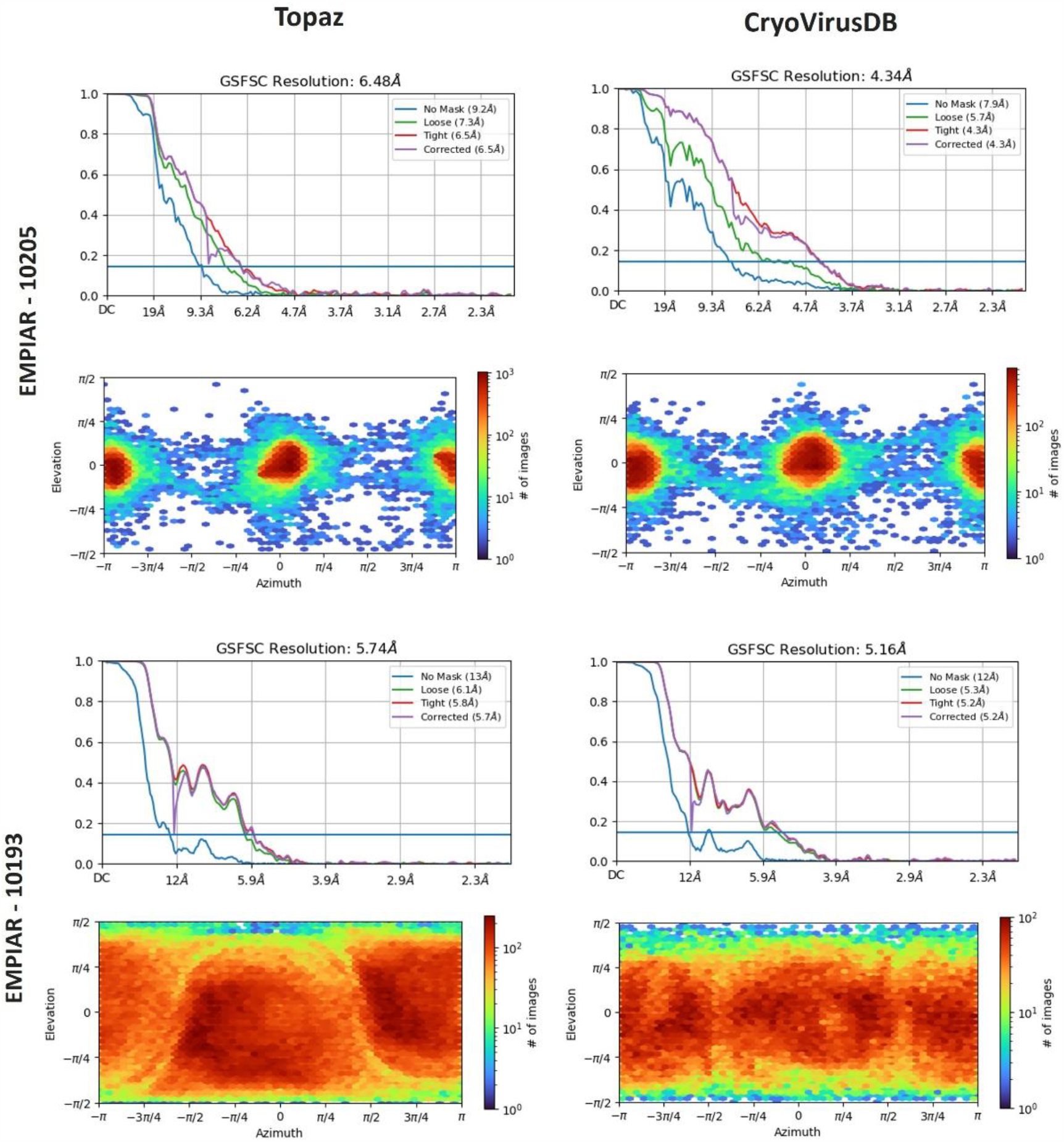
The comparison of 3D density resolution and direction distribution obtained by Topaz and CryoVirusDB on EMPIAR 10205 and EMPIAR 10193.

In the case of EMPIAR 10205, Topaz picked around 75,000 more particles compared to our manual picking. Despite this, the density map reconstructed from particles selected by CryoVirusDB achieved a resolution of 4.34 Å, substantially better than Topaz’s resolution of 6.48 Å. This indicates the high quality of the picked particles in CryoVirusDB. For EMPIAR 10193, the resolution of CryoVirusDB is 5.16 Å, also better than 5.74 Å of Topaz.

The heightened intensity of the red color in the direction distribution shown in the lower section of each block in **Figure 7** corresponds to an increased number of particles in the elevation vs azimuth plots. CryoVirusDB demonstrated superior particle picking by capturing a substantial number of particles with a wide angular distribution, evident in the red coloration on the heatmap for both validation cases.

The detailed comparison of the 3D density map reconstruction of three trials for CryoVirusDB and Topaz is provided in **Table 4**. The density maps constructed from the labeled particles in CryoVirusDB consistently exhibit a higher quality than Topaz in terms of multiple resolution metrics, even though the number of particles in CryoVirusDB is much smaller than the number of particles picked by Topaz, indicating that Topaz may pick quite some false positives and/or miss some true positives representing different views.

**Table 4:**
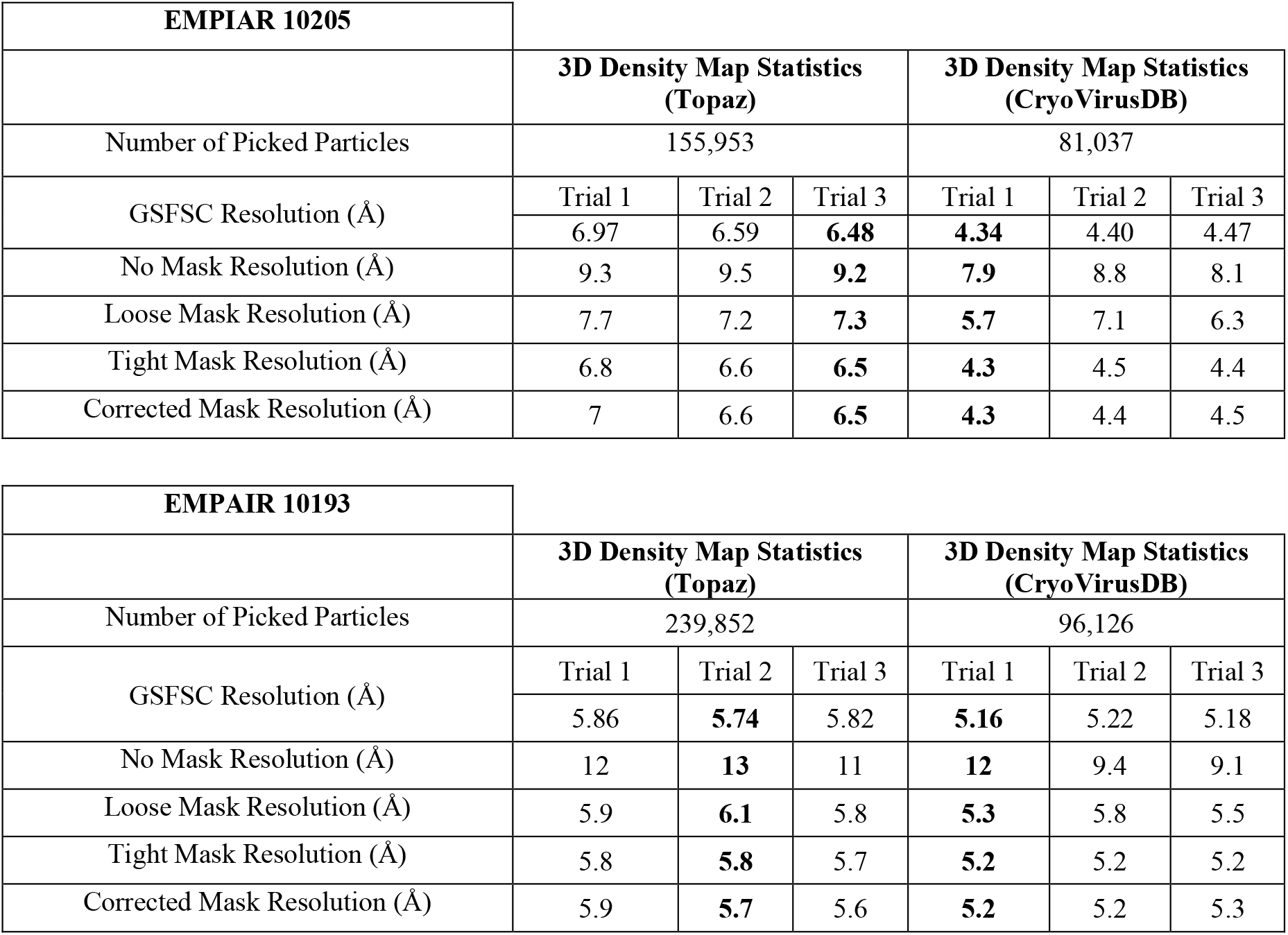
3D density map result comparison for EMPIAR 10205 and EMPIAR 10193. Bold fonts highlight the resolution of the best of the three trials for each method.

## Code Availability

The GitHub repository: https://github.com/BioinfoMachineLearning/CryoVirusDB contains all the scripts used in every stage of data curation. It also provides instructions on how to download and use the data.

## Author Contributions

J.C. conceived the research. R.G., A.D., and J.C. designed the methodology and experiment. R.G. and A.D. wrote the scripts and codes for dataset preprocessing. R.G. and A.D. curated the data. J.C. and L.W. conceptualized data validation. R.G. and A.D. drafted the manuscript. J.C and L.W. revised manuscript. All authors participated in result discussions, data analysis, and made contributions to the final manuscript.

## Competing Interests

The authors declare no competing interests.

## Funding

This work was supported by National Institute of Health grant (grant #: R01GM146340) to JC and LW.

## Notes

https://github.com/BioinfoMachineLearning/CryoVirusDB

### Competing Interest Statement

The authors have declared no competing interest.

